# Extending ligand efficacy indices with compound pharmacokinetic characteristics towards holistic Compound Quality Scores

**DOI:** 10.1101/2023.08.30.555339

**Authors:** Christofer S. Tautermann, Jens M. Borghardt, Roland Pfau, Matthias Zentgraf, Nils Weskamp, Achim Sauer

## Abstract

The suitability of a small molecule to become an oral drug is often assessed by simple physicochemical rules, the application of ligand efficacy scores (combining physicochemical properties with potency) or by multi-parameter composite scores based on physicochemical compound properties. These rules and scores are empirical and typically lack mechanistic background, such as information on pharmacokinetics (PK). We introduce a new type of Compound Quality Scores (specifically called dose-scores and c_max_-scores), which explicitly include predicted or when available experimentally determined PK parameters, such as volume of distribution, clearance and plasma protein binding. Combined with on-target potency, these scores are surrogates for an estimated dose or the corresponding c_max_. These Compound Quality Scores allow for prioritization of compounds in test cascades, and by integrating machine learning based potency and PK predictions, these scores allow prioritization for synthesis. We demonstrate the complementary and in most cases the superiority to existing efficiency metrics (such as ligand efficiency scores) by project examples.

## 1. Introduction

Oral drug delivery is the preferred administration route for small molecules mainly due to high convenience and compliance. A long history of failures and learnings [1] helped to define typical physicochemical ranges, for which a small molecule is expected to be orally bioavailable. Especially attrition in early clinical phases due to insufficient PK characteristics of drug candidates has been observed in the 1990s and early 2000s and could have been avoided by picking candidates with a more suitable PK profile. This led to a property-centric view in drug design, where most prominently Lipinski’s Rule of 5 (Ro5) [2] has been a hallmark in providing guidance on the the physicochemical properties of oral drugs. Specifically the molecular weight, the clogP and the hydrogen-bond donors and acceptors were set into relation with bioavailability [3] and are confined by upper thresholds. These empirical rules were derived from a set of back then marketed drugs and - although there are many exceptions - had a clear impact on the mindset of medicinal chemists in lead optimization (LO) and significantly limited the accessible chemical space in oral drug design. Twenty five years after the publication of the Ro5, it became clear that rule-based compound prioritization might be too simplistic [4, 5]. Many recently approved drugs break the Ro5, and average drug weight as well as lipophilicity have increased significantly over past decades, which might be a consequence of new target classes gaining attention [4]. To overcome the shortcoming of the Ro5, various new rules have been established, such as the Rule of 3 for fragment-approaches or others [6]. Important to note is, that all the rules were meant to confine the physicochemical property space to be compatible with oral drug administration, and such rules are used as more or less strict filters. Moreover rule-based approaches are not designed for ranking of compounds and, most remarkably, on-target potency is not even considered in these rules. However, drug design is a multi-parameter optimization, in which one strives for compounds with good potency on the target together with good PK properties as well as a clean off-target and toxicity profile - and this must be achieved in *one* single candidate. Therefore, rule-based approaches that emerged from analyses of the available drugs of the late 1990’s do not suffice to support innovative drug design today.

To overcome this strict rule-based paradigm, the idea of describing drug-likeness in terms of physicochemical properties has been further explored, leading to property scores, which describe the compound quality or “beauty”. The most common approaches are multi-parameter scores, which include the most prominent properties and condense these into one single score. Examples are the QED (quantitative estimate of drug-likeness) score by Bickerton et al. [7], in which the desirability of individual parameters is combined in one score and the approach by Wager et al., who designed a multi-parameter score specific for CNS-drugs based on empirical observations differentiating CNS-drugs from non-CNS drugs [8]. A more recent approach by DeGoey correlates preclinical PK results with cLogD, number of rotatable bonds, and number of aromatic rings [9]. All these approaches allow the ranking of compounds according to their drug-likeness and therefore are more versatile in application to rule-based approaches in this context.

Still, regardless of all physicochemical properties, a sufficient drug exposure at the target must be achieved in order to observe a desired physiological response. If the potency of a ligands is weak, higher compound concentration at the target side (or plasma concentrations as a surrogate) is required. Exceedingly high compound concentrations are at risk to cause undesired side effects due to off-target binding - and therefore, a high potency resulting in low efficacious drug concentrations is desirable. This fact is not covered by the rule-based approaches, but clearly triggered the next wave of compound scoring, where physicochemical properties are set in context with potency. Although this idea dates back to (at least) the 1980s, when Hansch phrased the guideline that drugs should be made as potent and hydrophilic as possible [10], the first modern approaches emerged in the early 2000s. Based on the observation that high molecular weight and a high clogP value are often detrimental to good PK properties (something also Lipinski referred to [3]), the potency of a ligand is set into relation with compound size and/or lipophilicity [11, 12, 13, 14]. This resulted in the ligand efficiency indices, which are based on various subtly different formulations. The most important are summarized in Table 1. The key idea of ligand efficiency indices is to estimate the price for higher potency in terms of molecular size and lipophilicity. Recent analyses of more than 3,000 chemical series showed that lipophilic ligand efficiency (LLE) tends to improve during optimization whereas ligand efficiency (LE) actually slightly decreases due to molecular growth [15]. This means that ligand efficiency indices can be considered powerful metrics for driving optimizations towards clinical candidates and can finally even be considered for candidate selection. In agreement, for most cases reported, candidates have clearly improved ligand efficiency indices (especially LLE) compared to the starting points (e.g., HTS hits) [16]. To test the usefulness of ligand efficiency indices in compound design (i.e., in compound ranking), the %-better metric in LO programs has been introduced [14, 17]. In this metric, the fraction of molecules with a *better* ligand efficiency than the final drug candidate is recorded - this can be applied for a LO campaign or to compare with a set of external molecules (e.g., from ChEMBL). In the examples provided [14, 18], only a small fraction shows better LE and LLE than the eventual drugs. Only for targets like kinases, where optimization is mostly driven by selectivity issues, ligand efficiency seems to be taken less into account during optimization.

**Table 1:**
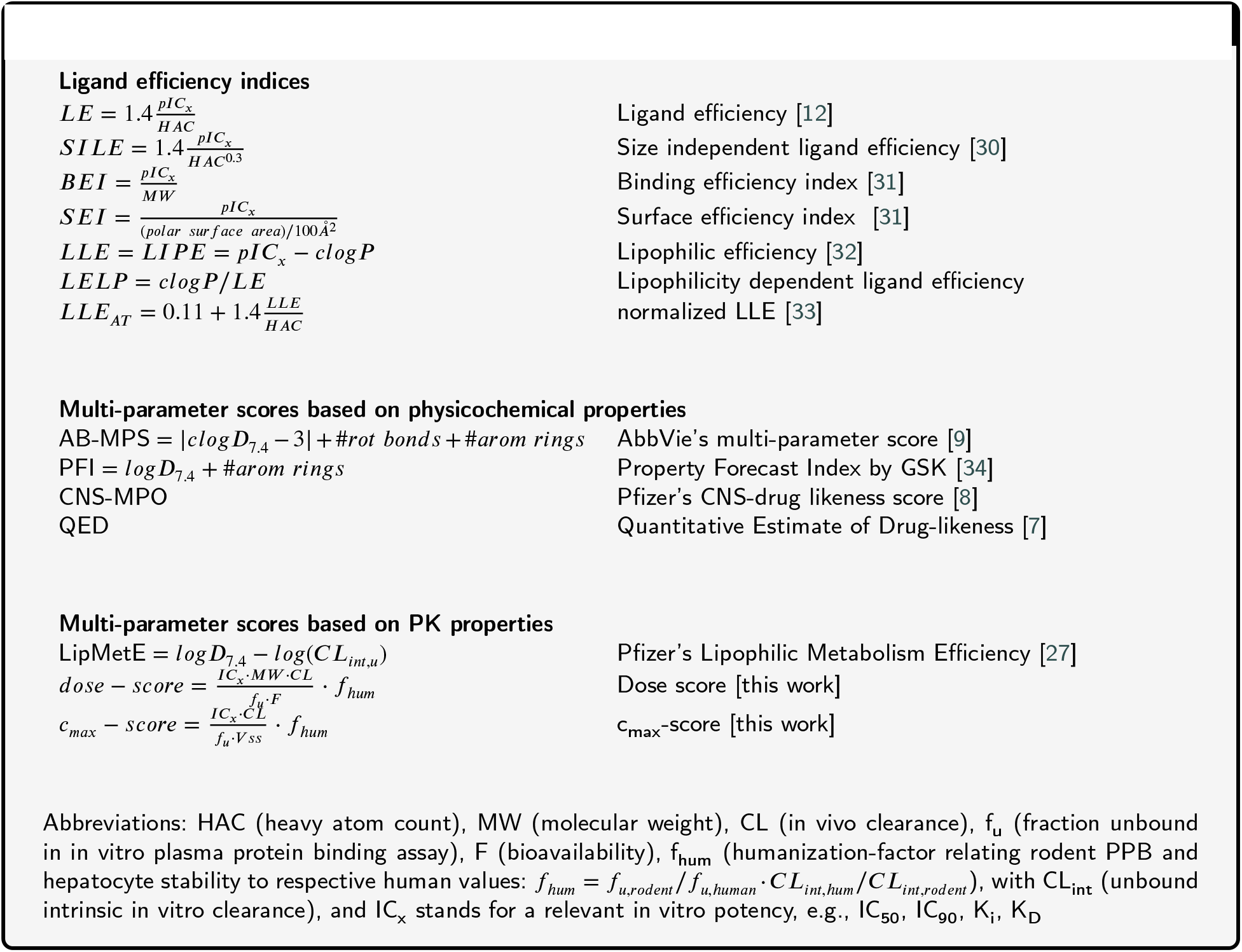
Ligand efficiency indices and multi-parameter scores.

Over the years, a large number of ligand efficiency indices have been proposed and can be put into numerical and algebraic relationship [19], as they are often based on common elements. Based on the multitude of available indices, there are also controversial discussions on the physical foundation [20, 21] as well as on the proper application [21, 22, 18]. Despite these fundamental discussions, many studies demonstrated the successful application of indices in compound optimization [22, 23, 17, 15, 18, 24, 16]. When looking at this from a different angle, efficiency indices are a pragmatic combination of potency and few physicochemical parameters. Eventually, lowering lipophilicity and mass provides only a very indirect link to PK properties and the need of a better and explicit inclusion of PK in compound optimization is emphasized. [25, 26] Indeed the idea of including PK-properties into efficiency scores recently emerged [27, 28] and the requirement for optimization towards estimated human doses has been underlined [29]. As a logical evolution it would be considered beneficial to combine both concepts in one score, i.e., to include the advantages of ligand efficiency indices, which normalize potency and combine this with compound-specific PK-properties. Therefore, we introduce the the *dose-score* and the *c*_*max*_*-score*, two multi-parameter scores derived from expressions estimating the oral dose required for efficacy and the corresponding maximal plasma concentration as a surrogate for safety. The advantage of the new scores is the explicit combination of PK-properties with potency resulting in multi-parameter scores that have a direct relevance for drug design.

## 2. Discussion: Multi-parameter scores based on PK properties

The goal of an LO campaign is to identify an efficacious and safe drug candidate by balancing a multitude of parameters. From a PK perspective the candidate has to reach sufficiently high target engagement (i.e., free compound concentration normalized by potency) to elicit the desired efficacy, while keeping the overall concentrations as low as possible to ensure safety. Therefore, we focus on two key parameters: the eventual efficacious oral dose and the corresponding c_max_. As we aim to develop a generic approach, we make some assumptions upfront. First, efficacy is achieved by unbound concentrations and second that a relevant in vitro potency value (e.g., IC_50_) has to be covered at C_trough_. Aiming for sufficiently high concentrations at trough is normally considered a conservative approach to predict the dose [29]. This implies that we do not account for potential hit-and-run mechanisms, and assume that a predictive in vitro potency assay is available. For dose estimation very thorough approaches are discussed [35], however, to keep it generic and tractable, we apply a simple one-compartment PK model to predict the concentrations and the respective required dose [36]. The detailed derivation is provided in the Appendix A. Several approximations were done, such as linearization of the exponential terms as well as neglecting various pre- and conversion factors. The resulting general form of the scores is:

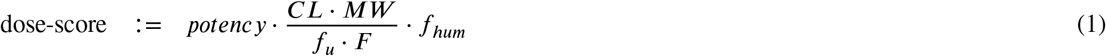

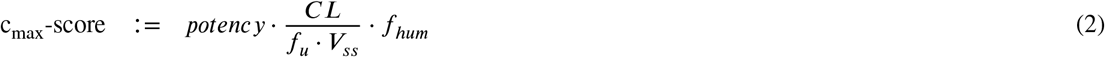

with the individual factors described in Appendix A as well as in Table 1. The structure of these scores, herein named “Compound Quality Scores”, strongly deviates from the known scores summarized in Table 1, for example no explicit lipophilicity terms are present. The equations above define a family of scores, depending on the in vivo PK species (here restricted to rat and mouse) and whether the “humanization” factor is applied. Overall, we discuss four different variants of the dose-score and the c_max_-score each (see Appendix A). The modular description allows for a straightforward extension to other species. Here, we focus on rodent in vivo PK values due to in the high amount of available rodent PK data (compared to higher non-rodent species).

So far, we have not discussed the origin of the PK parameters in Equations 1 and 2. Ideally these are measured experimentally, however in vivo data in early project phases is quite scarce. Still, we are propagating the application of the Compound Quality Scores very early in compound profiling (e.g., immediately after the primary on-target assay has been tested) or even during compound design. Therefore, measured PK parameters to calculate the Compound Quality Scores for the respective compound are rather the exception, and we use machine learning (ML) predictions trained on either public domain or in-house data sets to account for missing PK data [37, 38]. As a pragmatic approach, we replace these predicted PK parameters as soon as experimental data for a compound becomes available - usually in vitro before in vivo data in a project setting. This means that whenever new experimental results become available, they contribute to a more robust scoring.

As exemplified in Figure 1, the correlation of the PK parameters to the actual experimental values is better if some parameters have been experimentally determined. In a nutshell - the more experimental parameters known, the better. This is not given upfront, especially if parameters are highly correlated, error compensation may occur. In our experience, however, we see strong benefits by including experimental results. We shortly discuss the procedure to train our ML models and their prospective performance in project applications in Appendix B. As a positive example, plasma protein binding is predicted very well in a prospective setting with Spearman rank correlation coefficients above 0.8. Due to the increasing number of experimental values and new ML approaches, the prediction of non-clinical PK parameters by ML approaches has reached a satisfying level of precision [39]. However, complex endpoints, such as bioavailability are still challenging for prediction [40]. The scope of the present study, however, is not the discussion of the accuracy of predictions, but the introduction and the usage of the Compound Quality Scores allowing better guided compound optimization.

**Figure 1:**
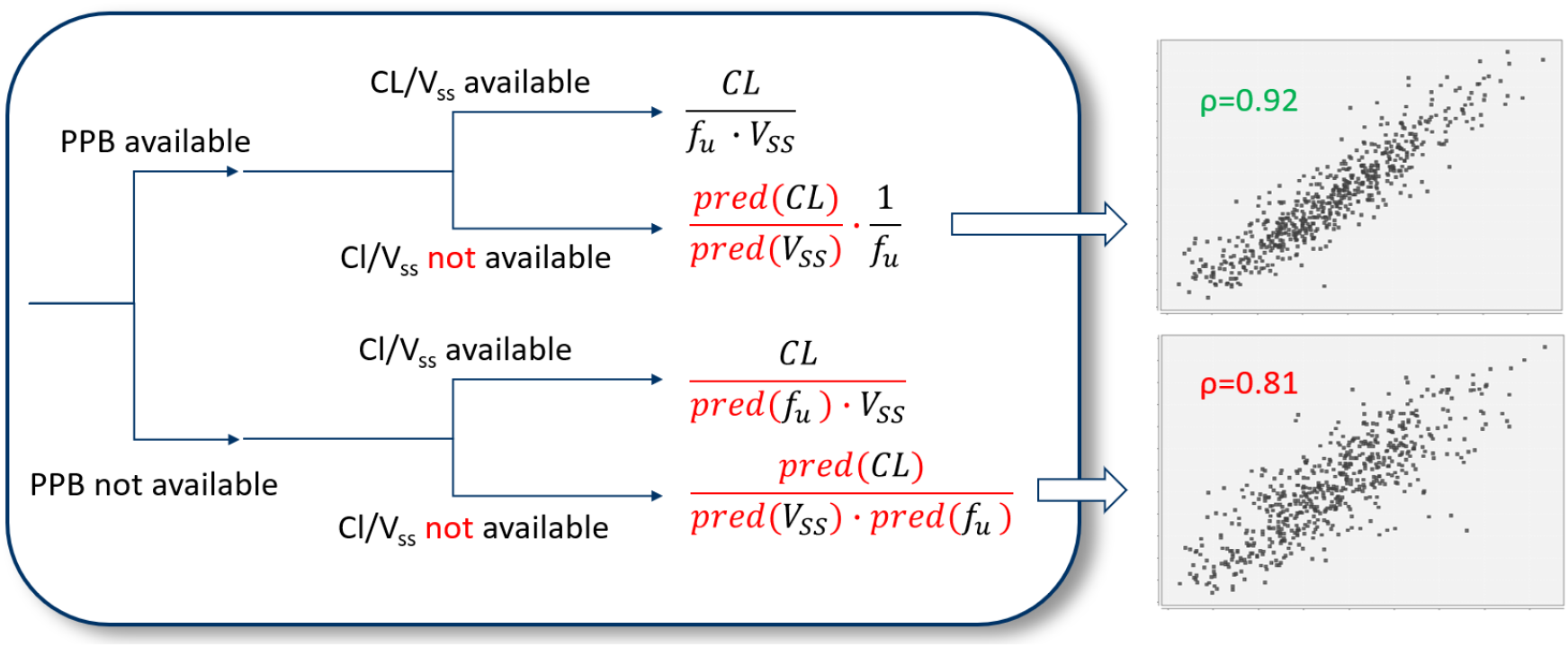
Description of the computation of the PK-component of the c_max_ -score 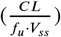. If all PK parameters are available from experiments, the experimental values are taken (upper leg), if no experimental PK parameters are available, all parameters are predicted in silico (*pred*({*CL, f*_*u*_, V_ss_}) (lowest leg). In case one or more, but not all PK parameters are available, experimental values are combined with predicted values to yield the PK-component of the score. On the right the impact of predictions on the accuracy of the PK-component is shown (log scale): correlation between the PK-component based on experimental data (x-axis) vs. the PK-component, where only *f*_*u*_ is measured (upper right, y-axis) or the PK-component that is solely based on in silico predictions if no experimental data is available (lower right, y-axis). Models were trained based on 4-fold cross validation [38] and *ρ* corresponds to the Spearman rank correlation coefficient.

### Score performance

To assess the applicability and potential impact of the Compound Quality Scores, we are comparing their performance with the current state of the art scores, which are LLE and LE for target-dependent metrics. In addition, we are illustrating how the scores have been applied in in-house optimization programs at Boehringer Ingelheim (BI) Pharma towards candidate selection.

Several studies demonstrate the power of LE and LLE in optimization programs [16, 14, 22, 17, 18, 15] and balancing between on-target potency and compound size and lipophilicity has proven to be a clear step forward in drug design compared to rule-based approaches [22, 17]. One way to demonstrate the applicability of scores and efficiency indices is to test whether approved drugs are correctly classified as the compounds with the highest scores. This test procedure is supported by two observations, first that approved drugs have higher LLE (and to a certain extent also higher LE) values compared to lead compounds or publicly known compounds for the corresponding target[14, 16, 18]. Second, a study by Perola [16] demonstrated that in a set of 60 lead/drug pairs LLE is higher for the drug in ∼80% of the examples. In line with this study, we considered the same 60 pairs and tested how often the actual drug provides better scores compared to the lead molecule. Table 3 in Appendix C displays the %-better metric [14, 17]. We observe that the newly introduced dose-scores and c_max_-scores perform much better compared to both LE and LLE. Especially LE was demonstrated not useful to prioritize compounds, a behavior that has recently also been shown for an even larger data set [15]. One may argue that the task of prioritizing a single lead over other drug molecules is not a real-life scenario and indeed, the actual task is to identify most promising compounds during LO to proceed with them more quickly. Therefore, the %-better metric is usually used to compare candidates and/or drugs with e.g., published compounds from sources like ChEMBL or all other compounds from an optimization campaign [17]. For this purpose, we choose ten discovery research projects at BI Pharma aiming at different targets, which all finally provided oral drug candidates that either entered or are about to enter clinical trials. To ensure diversity, this data set is comprised of four GPCRs projects (from various classes), one kinase, one serine protease, one ion channel, one other receptor and two enzymes, with a median of about 5,100 compounds per optimization program (compound inclusion criterion was the testing in the individual primary potency assay). Four of the programs were aimed to the CNS, thus transporter mediated efflux at the blood brain barrier has also been considered in our scores (see Appendix A). The performance of the Compound Quality Scores as well as the c_max_-scores is shown in Appendix C and in Figure 2. Very clearly, the newly defined scores outperform LE and LLE, which by themselves already impressively demonstrate their discriminating power between the (clinical) candidates and the rest of the program. While these metrics reveal the advantage of including PK parameters in multi-parameter scores, the application during compound design in LO projects is the typical and most important use case for medicinal chemists.

**Figure 2:**
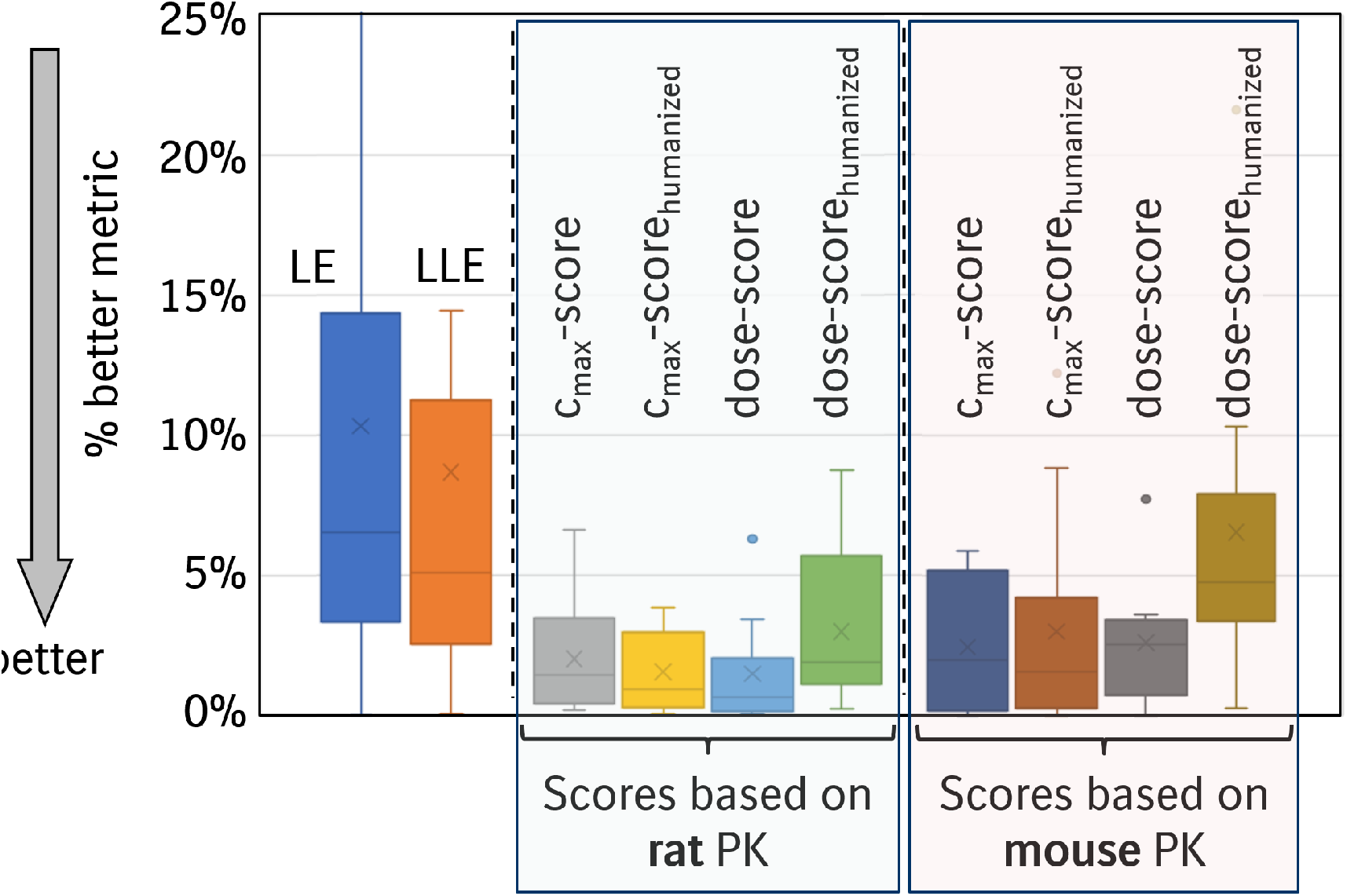
Comparison of the performance of LE and LLE scores compared to dose-scores as well as the cmax_max_-scores. The %-better metric describers the fraction of molecules in the individual optimization program that is actually better than the final (clinical) candidate. The lower the metric, the better is the score in prioritizing molecules with best profiles. The bars are derived from n=10 programs with more than 5,000 compounds on average.

### Use in optimization projects

When using the scores in project work, the choice of the most suitable score (out of the eight variants as shown in Figure 2) is not always straightforward. However, the Compound Quality Scores incorporate rodent in vivo data. Therefore, there is a canonical first choice to select the scores in optimization according to the pharmacodynamic (PD) species. In addition, we have observed, that it is often beneficial to have a close look at all 4 scores (dose-score, dose-score_humanized_, c_max_-score, and c_max_-score_humanized_) in the beginning of an optimization program, to decide which of the scores resembles manual rankings by experts (aiming for efficacy in PD models) the best. The more parameters that contribute to a score, the higher may be the accumulation of prediction errors and especially endpoints of higher complexity (such as bioavailability) will contribute quite a bit due to error propagation. Therefore, in our experience, the c_max_-score is the most robust one in terms of prediction errors, but the dose-score tends to be the score, that is closest to the way how project teams prioritize compounds. Clearly, here we see the benefit of including experimental rather than predicted parameters - the more experimental data included in the Compound Quality Scores the lower the uncertainties induced by prediction errors.

### Optimization Trajectories

Optimization trajectories are powerful in tracking the progress of a project [24] and we have picked a recent in-house GPCR project, where we demonstrate how the dose-score compares to efficiency metrics. In Figure 3 we show the progression of selected key compounds together with different parameters. Obviously, the first step of optimization was targeting to improve potency, which is visible in the strong improvement in IC_50_ and increase in LLE. However, in this project, the team also had a very close eye on additional compound properties, and in the first optimization phase the efflux scaled dose-score was decreased even more substantially, demonstrating, that the optimization was clearly directed towards an acceptable PK for CNS compounds. In the later stages of optimization, the dose-score could be kept in a favorable range towards delivering a candidate. Even though such optimization trajectories are very powerful to track optimization, as they are based on manually defined key compounds, their application is not easily implemented in compound design. For design it is more important to rank and to prioritize compounds accordingly, either for synthesis or for quick progression in the assay cascade towards in vivo assays.

**Figure 3:**
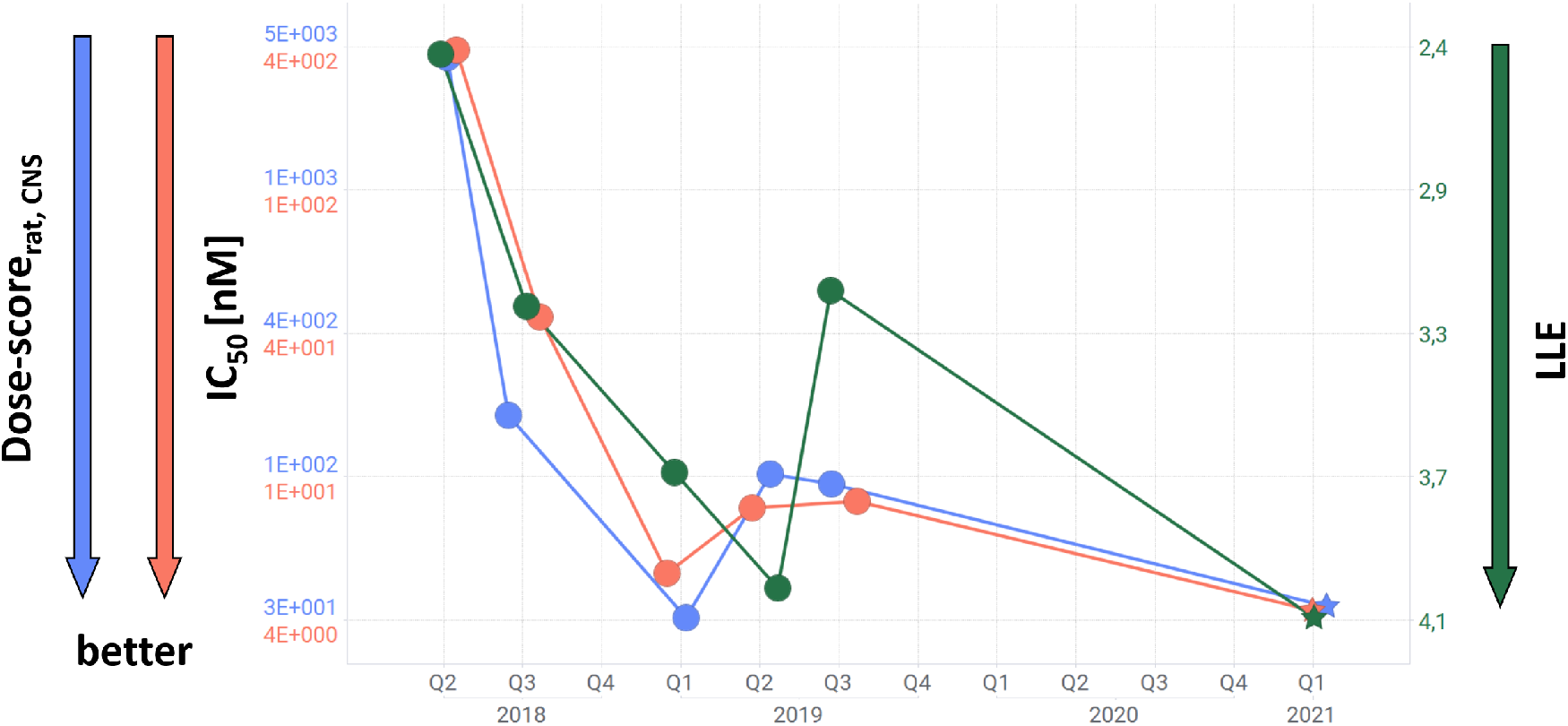
optimization time trajectory of a prototypical inhouse (GPCR) research project. Key compounds during optimization (circles) and the candidate (asterisk) are displayed and the corresponding LLE, potency and dose-scorerat,CNS values are displayed. LLE is depicted on a reverse axis to facilitate comparison.

### Ranking and prioritizing compounds

As demonstrated above, the Compound Quality Scores are well suited to identify the best compounds in a large data and in the time-trajectories we demonstrated their ability to illustrate the progress of a project. Now, we combine these properties and provide a more detailed opportunity to prioritize compounds. Figure 4 was generated in the same GPCR project as Figure 3, but now all compounds with a potency below 9μM are displayed. Compounds, for which non-rodent PK studies have been performed, are assumed to be most promising compounds in a project, as they guide the path towards the final candidate and these compounds are highlighted in Figure 4 in green. In subplot (A), the dose-score_rat,CNS_ is plotted against the IC_50_. Ligands with the same potency show up at the same x-axis value, but the difference in their y-axis values corresponds to the spread in the quality of their PK-properties. The lower the dose-score, the better the PK quality so that in general, the lower rim of the cloud contains the most important compounds, which should be moved into the next stage of the screening cascade, until a showstopper (such as selectivity, toxicity) is identified. In agreement with that, the compounds selected by the project team to perform non-rodent PK studies are mostly located on the lower part of the cloud in Figure 4(A). In comparison, we display the same compounds in Figure 4(B) with the corresponding LLE. We clearly see, that in this example the guidance by LLE for compound prioritization is less powerful, as the location of the non-rodent PK compounds is somewhat in the indistinct middle of the cloud.

**Figure 4:**
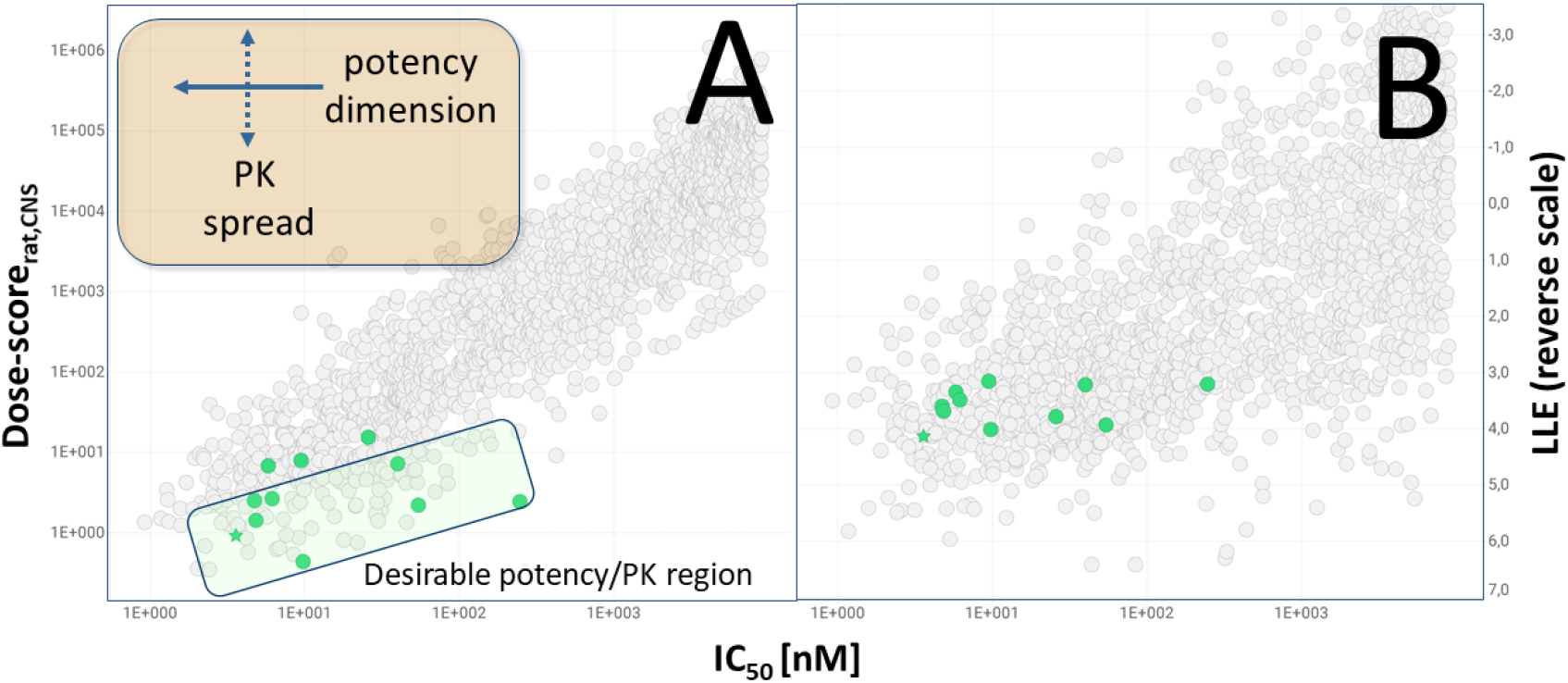
Plots of the IC_50_ vs. dose score (A) and LLE (B) from an in-house GPCR project. Both plots display the same set of about 1700 compounds with a potency better than 9 μM. Compounds colored in green have been tested in non-rodent species PK and therefore had a high importance to the project. The final candidate is denoted by a green asterisk.

## 3. Conclusions

We have introduced the Compound Quality Scores (dose-score and c_max_ -score), that combine potency as well as PK -parameters into one multi-parameter score. These scores are designed so that they allow the ranking of compounds according to an approximate dose and approximate c_max_ value based on simple assumptions regarding the PK/PD relationship (C_through_-driven efficacy). Applying additional approximations of basic PK formulae led to very simple scores that are useful for ranking and prioritizing compounds. However, due to the simplifications the scores are not intended to yield absolute values for efficacious doses and are not intended to be used for cross-project comparisons. We demonstrate the superiority of the new scores over state of the art metrics, such as LLE, in the identification of drug candidates in a drug discovery project. We also demonstrate the usefulness in compound ranking for lead optimization purposes. Metrics like LLE and LE include PK only implicitly through the assumption that high clogP and MW are inversely correlated to good PK properties. The large difference to all these existing scores so far is the explicit inclusion of PK parameters together with the potency, yielding a single score for the compound quality in terms of ADME and efficacy. If no experimental PK parameters are available, predictions from machine learning models can be used for single PK endpoints, still allowing for acceptable to good accuracy. In the course of a project, once more experimental parameters become available, the scores become more reliable (i.e., less prediction-error prone) over time. Therefore, we consider that our approach, despite all its approximations and possible prediction errors is a step forward in the generation of meaningful multi-parameter-scores and we recommend to add the Compound Quality Scores into the repertoire of metrics that are regularly used for medicinal chemistry optimization purposes.

### A. Derivation of dose-scores c_max_-scores

We use the simplest equations based on a one-compartment PK model with instantaneous absorption for the required dose [36] and the plasma concentration after dosage

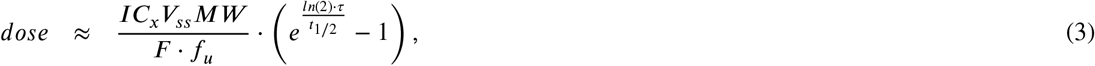

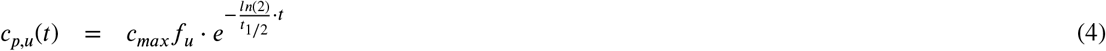

with

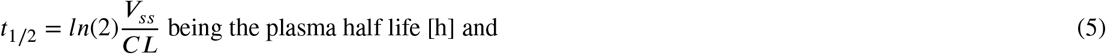

V_ss_ [l/kg] is the volume of distribution at steady state, F [%] is the bioavailability, CL [l/h/kg] is the in vivo plasma clearance after i.v. administration, MW [g/mol] is the molecular weight, f_u_ [%] is the fraction unbound, c_max_ [M] is the maximal total plasma concentration after single dosage and c_p,u_(t) [M] is the total unbound plasma concentration at the time-point t [h] after dosing. IC_x_ [M] is the surrogate for the desired trough target coverage (e.g. IC_50_ for a through inhibition of at least 50%) for the desired dosing interval *τ* of 12 or 24h. Equation 3 is a simple approximation for through coverage of a IC_x_ concentration for a single *oral* administration, whereas Equation 4 is a description of a single compartment PK model (i.e., a monophasic / monoexponential PK profile) after having reached the maximal concentration independent of administration. With these two equations we can formulate the goals for compound optimization in terms of PK properties, assuming that the pharmacodynamic (PD) effect requires a through coverage of IC_x_. Equation 3 directly allows to formulate the goal *dose* → *min* and when inserting IC_x_ as lower bound for c_p,u_, equation 4 rewrites to

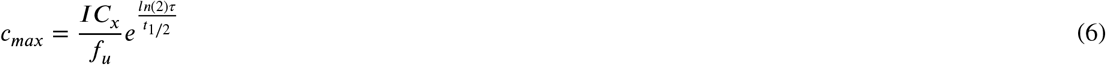

and the goal is to optimize candidates with low total maximal plasma concentrations (in other words, with a low peak/through ratio) to avoid off-target activity.

#### A.1. Linearization and Score formulation

With the optimization parameters (low dose and low c _max_) as defined above, we now aim to define tractable scores, that are useful for guiding compound optimization. Equations 3 and 6 both do have exponential terms and are therefore very sensitive to uncertainties in the exponents - this is an unwanted behavior because eventually the scores will be fed by machine learning (ML) predictions and should be tolerant to a small degree of uncertainty (which can also be present in experimental in vitro or in vivo data). We therefore decided to linearize Equations 3 and 6 and truncation of a Taylor expansion of the respective equations leads to

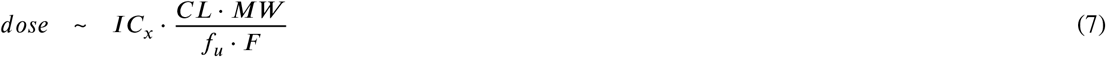

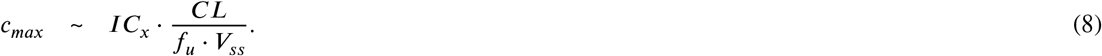

To come to these equations, *τ* is assumed to be arbitrary but fixed (e.g., 12h or 24h), and the Taylor expansion is done over CL at the point 0 (so actually being a Maclaurin expansion). Truncation after the linear factor and omitting all resulting constant pre-factors (e.g., coming from conversions between units) yield Equations 7 and 8 (“∼” rather than a “=” indicates the simplifications). We do not aim to develop scores for inter-project comparison, but for ranking of compounds within a project during optimization, therefore we are not interested in the pre-factors . An equivalent formulation of Equation 7 has been deduced by Gunaydin et al. [36] by alternative considerations.

#### A.2. Species and score humanization

So far, in the derivation of the formulae we have not discussed the species for the PK data and implicitly have assumed to work with human data. This is unfortunately a too optimistic assumption as in all practical applications usually rodent PK data is available to guide compound optimization. On one hand we need parameters like CL, Vss and F, where human data is very scarce and we need a way to bridge to rodent data. On the other hand, we can quite easily get the parameters for human IC_x_ and human f_u_ from in vitro assays. This allows now for several advanced formulations of the scores reflecting the human situation at least in part. The most straightforward way is to solely rely on rodent PK data and only use human IC_x_ from in vitro assays, as shown in the definitions in Equations 9 and 10. The advantage of Equations 9 and 10 is that for most of the parameters a wealth of data exists in the public domain as well as in company proprietary databases and quite decent ML models can be trained for compounds where experimental data is missing.

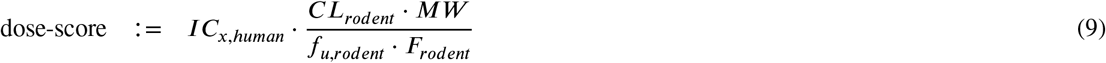

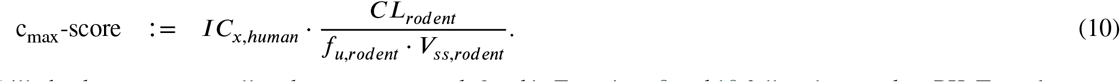

Still, the dose-score as well as the c_max_-score as defined in Equations 9 and 10 fully rely on rodent PK. To make a step towards more “humanized” scores, we take the approach to first, include human plasma protein binding data and second, scale the rodent in vivo clearance by the human/rodent ratio of clearance observed in hepatocyte assays in vitro. For and bioavailability we decided that the PK parameter identified in rodents is assumed also for the human situation. is sufficiently well conserved over species [29] and staying with V is not the most strongest assumption in definition of the scores. And here we need again to emphasize that scores are *not* meant to give quantitative estimates of human doses, but they are designed for compound ranking. systematic error in V_ss_ within a compound class would not at all impact the ranking and, thus we go forwards with V_ss,rodent_. A similar line of argumentation holds for bioavailability, although the linear correlation between F_human_ and F_rodent_ is weak. However, classification in high and low bioavailability seems to be feasible though [41]. With this we define the humanized scores as

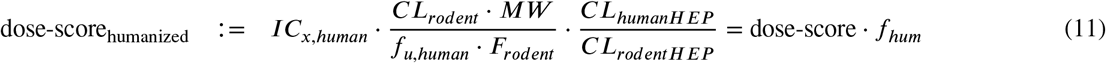

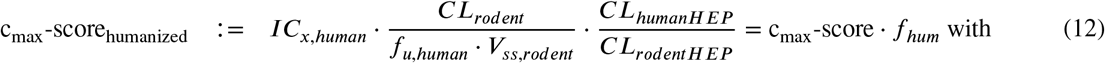

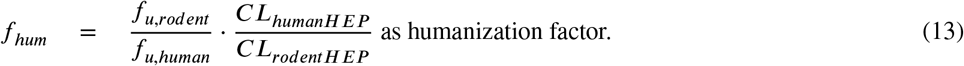

CL_humanHEP_ and CL_rodentHEP_ correspond to the intrinsic clearance in human and rodent hepatocyte assays in vitro performed under comparable conditions. Both scores, dose-score_humanized_ as well as c_max_-score_humanized_ can be seen as the extension of the rodent counterparts dose-score and c_max_-score multiplied with the humanization factor *f*_*hum*_ from Equation 13. The humanized scores are obviously more relevant for optimization, as the goal is to deliver drugs for humans. However, the reason why we discuss also the rodent scores comes from the fact that they rely on less parameters and thereby introduce less potential errors due to inaccurate predictions. There are also approaches available, that directly predict human PK parameters, but they are based on a much smaller data basis [42, 43] and don’t allow for model improvement by additional experimental in vivo data during LO.

### Special case: CNS drugs

Drugs which exert their action in the central nervous system (CNS), have to cross the blood brain barrier in order to reach their target. One decisive parameter, that describes the blood-plasma vs. brain concentrations is the efflux ratio at the blood brain barrier. Most important in this context is the efflux transporter P-glycoprotein (P-gp). The efflux ratio is derived from in vitro experiments in MDCK cells overexpressing P-gp, which correlates well with the in vivo efflux at the blood brain barrier of rodents [44] To include the case of CNS drugs, we therefore need to include the efflux-ratio into the definition of our scores, i.e.,

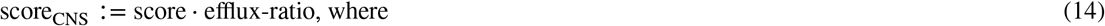

“score” stands as placeholder for any score defined in Equations 9-12.

### B. Prediction of PK endpoints

**Table 2:** Statistics for predicted PK-parameters in research project work at BI in the time-range of January 2021 - August 2022. Machine learning models have been re-trained every 2 months and the predictions in this table are purely prospective. For the first 8 rows, the statistics are based on log_10_ scaled values and for the bottom 5 rows the statistics are based on non-scaled values. # examples refer to experimental data points, which were determined over the provided time range. Training of regression models has been described elsewhere [38]. Specifically, here we employ extremely randomized trees [45] in the context of random forest regression with 600 estimators as implemented in scikit-learn [46] based on 4,414 real-valued descriptors [47].

| PK endpoint | Pearson $r^2$ | Spearman $\rho$ | RMSE | # examples | |
| --- | --- | --- | --- | --- | --- |
| MDCK efflux ratio | 0.62 | 0.78 | 0.39 | 3541 | log-scale |
| $f_{u, \text{rat}}$ | 0.68 | 0.83 | 0.48 | 1659 | |
| $f_{u, \text{mouse}}$ | 0.71 | 0.84 | 0.47 | 1425 | |
| $f_{u, \text{human}}$ | 0.70 | 0.84 | 0.47 | 3202 | |
| $V_{ss, \text{rat}}$ | 0.35 | 0.53 | 0.36 | 542 | |
| $CL_{\text{rat}}$ | 0.32 | 0.39 | 0.47 | 475 | |
| $V_{ss, \text{mouse}}$ | 0.32 | 0.59 | 0.35 | 450 | |
| $CL_{\text{mouse}}$ | 0.34 | 0.57 | 0.42 | 443 | |
| %Qh HEP<sub>rat</sub> | 0.44 | 0.66 | 21.9 | 978 | not scaled |
| %Qh HEP<sub>mouse</sub> | 0.40 | 0.63 | 22.1 | 1092 |  |
| %Qh HEP<sub>human</sub> | 0.42 | 0.62 | 19.6 | 2550 |  |
| $F_{\text{rat}}$ | 0.10 | 0.33 | 33.9 | 132 | |
| $F_{\text{mouse}}$ | 0.22 | 0.45 | 29.7 | 302 | |

### C. %-better metrics

**Table 3:** Comparison of the performance of LE and LLE scores compared to dose-scores as well as the cmax_max_-scores by the %-better metric. See the text for more details.

| Scores | LO program median (n=10) | Perola Leads/Drugs[16] |  |
| --- | --- | --- | --- |
| LE | 6.5% | 45.0% |  |
| LLE | 5.1% | 23.3% |  |
| dose-score | 0.6% | 13.3% | rat PK |
| dose-score<sub>humanized</sub> | 1.9% | 8.3% |  |
| c<sub>max</sub>-score | 1.4% | 8.3% |  |
| c<sub>max</sub>-score<sub>humanized</sub> | 0.9% | 6.7% |  |
| dose-score | 2.5% | 11.7% | mouse PK |
| dose-score<sub>humanized</sub> | 4.8% | 8.3% |  |
| c<sub>max</sub>-score | 2.0% | 8.3% |  |
| c<sub>max</sub>-score<sub>humanized</sub> | 1.5% | 5.0% |  |

